# Relationships and genome evolution of polyploid *Salix* species revealed by RAD sequencing data

**DOI:** 10.1101/864504

**Authors:** Natascha D. Wagner, Li He, Elvira Hörandl

## Abstract

Despite the general progress in using next generation sequencing techniques for evolutionary research questions, the analysis of polyploid species is still hampered by the lack of suitable analytical tools and the statistical difficulties of dealing with more than two alleles per locus. Polyploidization and especially allopolyploidy leads to new combinations of traits by combining genomes of two or more parental species. This enhances the adaptive potential and often results in speciation. However, multiple origins of polyploids, backcrossing to the parental species and post-origin evolution can strongly influence the genome composition of polyploid species. Here, we used RAD sequencing, which revealed 23,393 loci and 320,010 high quality SNPs, to analyze the relationships and origin of seven polyploid species of the diverse genus *Salix* by utilizing a phylogenomic and a network approach, as well as analyzing the genetic structure and composition of the polyploid genome in comparison to putative parental species. We adapted the SNiPloid pipeline that was originally developed to analyse SNP composition of recently established allotetraploid crop lineages to RAD sequencing data by using concatenated RAD loci as reference. Our results revealed a well-resolved phylogeny of 35 species of Eurasian shrub willows (*Salix* subg. *Chamaetia/Vetrix*), including 28 diploid and 7 polyploid species. Polyploidization in willows appears to be predominantly connected to hybridization, i.e. to an allopolyploid origin of species. More ancient allopolyploidization events involving hybridization of more distantly related, ancestral lineages were observed for two hexaploid and one octoploid species. Our data suggested a more recent allopolyploid origin for the included tetraploids within the major subclades and identified putative parental taxa that appear to be plausible in the context of geographical, morphological and ecological patterns. SNiPloid and HyDe analyses disentangled the different genomic signatures resulting from hybrid origin, backcrossing, and secondary post-origin evolution in the polyploid species. All tetraploids showed a considerable post-origin, species-specific proportion of SNPs. The amount of extant hybridization appears to be related to the degree of geographical and ecological isolation of species. Our data demonstrate that high-quality RAD sequencing data are suitable and highly informative for the analysis of the origin and relationships of polyploid species. The combination of the traditional tools RAxML, STRUCTURE, SplitsTree and recently developed programs like SNAPP, HyDe and SNiPloid established a bioinformatic pipeline for unraveling the complexity of polyploid genomes.

## Introduction

The doubling of genomes had a strong impact on diversification and evolution of complexity of several eukaryotic lineages (Van De Peer et al. 2009). Several studies have shown that polyploidization had strongly influenced the evolution of flowering plants (Soltis and Soltis 2009; Jiao et al. 2011). Approximately 25% of plants have experienced recent polyploidization and about 70% show signs of ancient genome duplication (Barker et al. 2016). Polyploidization creates “genomic novelty” and enhanced genetic diversity, thus increasing the ecological potential (Soltis and Soltis 2009). It is estimated that at least 15% of speciation events in angiosperms result from polyploidy (Wood et al. 2009; Mayrose et al. 2011). In general, two types of polyploidy are distinguished, based on the mode of origin: autopolyploidy and allopolyploidy. Autopolyploidy originates by genome duplication within the same species, whereas allopolyploids originate through interspecific hybridization and genome duplication, thereby combining duplicated sets of chromosomes from different parental species (Comai 2005). Autopolyploids are often hampered by irregular pairing of chromosomes at meiosis and often show multisomic inheritance (Comai 2005), thus autopolyploid cytotypes rarely evolve into distinct species (Soltis et al. 2007). Allopolyploidy allows for a more regular bivalent pairing and recombination of homologs (chromosomes from the same parent), whereas homeologs (chromosomes from the different parents) do not pair, resulting in a predominantly bisomic inheritance and fixed heterozygosity of the parental genomes (Comai 2005). Allopolyploids frequently exhibit novel morphological and ecological features, and hence are more readily recognized as distinct species (Soltis and Soltis 2009).

Although the impact of polyploidy in plant evolution is evident, the potential offered by the rise of NGS tools to analyze the origin of natural polyploid plant species is not fully exploited. The main reasons might be the difficulties in SNP calling, especially the need to distinguish different alleles derived from the same parent (homologs) from alleles originating from different parents (homeologs; Clevenger et al. 2015). Other difficulties are the necessity to filter paralogs (Hirsch and Robin Buell 2013), the uncertainty of allelic dosage as well as the lack of appropriate bioinformatic tools and models (Meirmans et al. 2018). Autotetraploid species possess nearly identical subgenomes, while allotetraploid species combine the two genomes of each parental species, each with two alleles. Given a sufficient coverage, NGS methods that generate hundred thousands of short reads are generally suited to detect all different alleles at a specific locus. However, merged sequences, as they are observed in most NGS pipelines, will lead to inaccurate results or weird positions of the allotetraploid in the tree. To face this problem, allele phasing can be used to analyze the origin and position of the tetraploid species in a phylogeny. Subsequently these data can be used for gene tree and species tree reconstruction. The separated (phased) homeolog alleles of an allotetraploid will group with the specific parental species (Eriksson et al. 2018). However, phasing tools assume diploid species, and thus, phased “alleles” in polyploids might contain allelic variants. The sorting of these variants could be done manually, as for example shown in Eriksson et al. (2018), but this is only feasible for a small number of loci, while NGS tools in general generate thousands of loci. Clevenger et al. (2018) used haplotype based genotyping for high-accurate SNP calling of polyploid subgenomes. However, both methods require long fragments and/or a reference genome. To use short read data to assemble and phase alleles or use haplotype-based genotyping for the analysis of polyploids is still challenging, especially, if no reference genome is available.

RAD sequencing (Baird et al. 2008) is a widely used reduced representation library (RRL) technique that harvest thousands of informative loci and ten thousands SNPs representing the whole genome without the need of previous genomic knowledge or a reference genome. This makes RAD sequencing a frequently used tool, especially for non-model organisms. The resulting variable sites can be used to analyse evolutionary processes. Furthermore, the thousands of biallelic SNPs harvested by RAD sequencing are generally well suited to resolve reticulate relationships, reconstruct species trees and to analyse polyploid species in a phylogenetic scope (Bryant et al. 2012; Dufresne et al. 2014; Clevenger et al. 2015). However, there are only few studies using reduced representation libraries like RAD sequencing data, when dealing with polyploid non-model plants (Chen et al. 2013; Mastretta-Yanes et al. 2014; Qi et al. 2015; Brandrud et al. 2019). While many independent loci are advantageous for the reconstruction of the polyploid origin, phasing of many unlinked short loci as generated by RAD sequencing will not reveal enough information if treated separately. A concatenation of the phased loci will reveal a chimeric mixture of alleles. Moreover, the tools to discriminate homeologs and their allelic variants in polyploids (more than two alleles) are not developed yet. To avoid these issues, a consensus sequence of all possible alleles can be used. The underlying allelic information is summarized in the consensus sequence in form of ambiguous sites at heterozygous positions. The resulting sequence information and SNPs will be used for downstream analysis. The common practice of neglecting allelic information for phylogenetic reconstruction has been criticized (e.g. Anderman et al. 2019) and is of course a specific problem in polyploids. However, these effects can be minimized by treating the polyploid species as “diploids” and reducing the allelic variants to two ‘alleles’=subgenomes. The calling of biallelic SNPs based on likelihoods can overcome biases associated with genotype uncertainty (Blischak et al. 2018b). Recently, Brandrud et al. (2019) presented a workflow based on SNP data to deal with polyploids in a recently evolved orchid species complex (*Dactylorhiza*). It points out the ability of SNPs to analyse polyploid taxa. Their workflow, however, is specifically suited to a closely related species group, but not to more diverged taxa.

The genus *Salix* L. (Salicaceae) comprises about 400-450 species of trees and shrubs that are mainly distributed in the Northern Hemisphere. In contrast to many other woody plant lineages about 40% of the species are polyploid (Suda and Argus 1968), ranging from diploid to octoploid species (rarely deca-/dodecaploid), based on the chromosome number *x* = 19 (Argus 1997). Molecular clock analyses suggested that the genus *Salix* originated in the early Eocene and started to diversify in the middle Eocene (Wu et al. 2015). Previous studies proposed that ancient genome duplications occurred several times in *Salix* (Dorn 1976; Leskinen and Alström-Rapaport 1999) and in the Salicoid lineage (Tuskan et al. 2006). Paleopolyploid species are more difficult to recognize, because their cytological behaviour has undergone ‘diploidization’, i.e. they have returned to regular chromosome pairing at meiosis as typical for diploid sexual species (Comai 2005). Therefore, polyploidization events can be best understood in the phylogenetic framework of diploid congeneric species. Recent allopolyploidization (young polyploidy) may have happened during the Pleistocene, when climatic oscillations caused range fluctuations of plant species and enhanced secondary contact hybridization of previously isolated species (Hewitt 2004; Abbott et al. 2013; Kadereit 2015). While homoploid hybridization may result in introgressive hybridization with the parental species (Hardig et al. 2010; Gramlich and Hörandl 2016; Gramlich et al. 2018), allopolyploidy can establish a rapid crossing barrier against the parents and hence may result in the evolution of an independent lineage. Cold climatic regimes during the Pleistocene, however, may have also triggered spontaneous autopolyploidization by unreduced gamete formation (Ramsey and Schemske 1998). These processes might have affected mainly willow species occurring in the temperate to arctic zones of the Northern Hemisphere, where glaciations had the most severe impact. In willows, natural hybridization is a frequent phenomenon (Argus 1997; Skvortsov 1999; Hörandl et al. 2012) and may occur even between distantly related species (e.g. Hardig et al. 2000; Gramlich et al. 2018). However, these processes are not necessarily exclusive. Hence, we expect to trace signatures of ancient and/or recent hybrid origin, secondary lineage-specific evolution, backcrossing and introgression in the genomes of willows. While in the 19^th^ century experimental crosses in *Salix* were popular and even triple or multiple hybrids have been produced even between distinct taxa (Wichura 1865), the origin of most natural polyploid species is still unknown. Thus, the genus *Salix* is an interesting system to study the different scenarios leading to polyploidy. About two thirds of the species occur in Eurasia, while about 140 species occur in America. Only a few species are native to Africa. Classical taxonomy and systematics in *Salix* have proven to be extremely difficult because of dioecy, simple, reduced flowers, common natural formation of hybrids, high intraspecific phenotypic variation and the presence of polyploid species (Skvortsov 1999; Hörandl et al. 2012; Cronk et al. 2015). However, related species are usually separated by distinct geographical distributions and/or ecological niches, often following altitudinal differentiation (Martini and Paiero 1988; Skvortsov, 1999; Hörandl et al. 2012). Phylogenetic analyses using traditional Sanger sequencing separated the lowland tree species (subgenus *Salix*) from the mostly shrubby species of a big clade uniting the two subgenera *Chamaetia* and *Vetrix*, but lacked any infrageneric resolution (Wu et al. 2015). The diversification of this clade started probably in the late Oligocene (Wu et al. 2015). We are mainly interested in the evolution of these shrub willows, which range from creeping arctic-alpine dwarf shrubs to medium-sized trees (Wagner et al. 2018). In the *Chamaetia/Vetrix* clade, which comprises about three quarters of all *Salix* species, more than one third of all species with known ploidy levels are tetra-, hexa- or octoploids. An analytical toolkit for the discrimination of polyploid willows has been established, based on the combination of SSR markers and flow cytometry, but was only used to detect the ploidy level for breeding purposes (Guo et al. 2016). In subgenus *Salix*, the tetraploid *S. alba* – *S. fragilis* complex has been analyzed with AFLP markers and revealed an allotetraploid nature of the species (Barcaccia et al. 2014), but in the *Chamaetia/Vetrix* clade the polyploid species have never been analyzed with molecular markers.

So far, studying the phylogenetic relationships in *Salix* was problematic because of a lack of resolution of nuclear and plastid based phylogenies (Percy et al. 2014; Wu et al. 2015). This lack of a phylogenetic framework for willows hampered also the reconstruction of relationships of polyploids within the genus. RAD sequencing (Baird et al. 2008) was recently used to overcome this lack of information in genus *Salix* (Gramlich et al. 2018; Wagner et al. 2018). This technique reveals thousands of genome wide informative SNPs to analyze relationships within and among species (Baird et al. 2008; Wagner et al. 2018). Wagner et al. (2018) published the first well-resolved phylogeny of diploid European members of the *Chamaetia*/*Vetrix* clade, while Gramlich et al. (2018) successfully used RAD sequencing to analyse recent homoploid hybridization patterns and introgression between two *Salix* species on alpine glacier forefields. RAD sequencing loci in *Salix* represent almost exclusively the nuclear genome, and provide ten thousands of SNPs from both conservative and rapidly evolving non-coding genomic regions (Gramlich et al. 2018). Therefore, this method is well suited to resolve reticulate relationships and to analyse the origin of the polyploid species in *Salix*.

In this study we want to test the utility of RAD sequencing data for the analysis of polyploid species with an emphasis on allotetraploids in an empirical dataset of diploid and polyploid shrub willows. We will use a comprehensive approach of different traditional and new analytic tools to elucidate different evolutionary scenarios of polyploid origin in Eurasian willows of the *Chamaetia/Vetrix* clade based on RAD sequencing data. Based on a well-resolved phylogeny, we want to test for the monophyly of the polyploid lineages. By using various approaches we will further test for (1) recent (Pleistocene) allopolyploidization by hybridization of two (or more) diploid parents; (2) more ancient (pre-Pleistocene) allopolyploidization of progenitors of extant species, and (3) autopolyploidization. Based on this information, we (4) try to get insights into post-origin evolution of polyploid species by analyzing their genomic composition.

## Material and Methods

### Sampling

For this study, we sampled 28 diploid species, one triploid, three tetraploid, two hexaploid and one octoploid species representing 19 sections sensu Skvortsov (1999) of the *Chamaetia*/*Vetrix* clade. The accessions of *Salix repens* ssp. *rosmarinifolia* and *S. lapponum* ssp. *ceretana* were treated as subspecies here. *Salix triandra* (subg. *Salix*) was included to serve as outgroup as closest relative to *Chamaetia/Vetrix* following the results of Wu et al. (2015). Hence, the complete sample set consists of 36 *Salix* species. The samples were collected in Central and Northern Europe as well as in China and determined after Fang et al. (1999), Skvortsov (1999) and Hörandl et al. (2012). The diploid species represent all main European lineages (Rechinger and Akeroyd 1993). Leaves were dried in silica gel and herbarium voucher specimens were deposited in the herbarium of the University of Goettingen (GOET). Two to five accessions per species were included in the analyses given a total of 133 samples. Detailed information about the sampling is summarized in Supplement Table S1.

### Ploidy determination using Flow Cytometry

To determine the ploidy levels of seven species without reported chromosome counts we performed flow cytometric measurements. The ploidy levels of 23 samples were estimated using flow cytometric analyses based on DAPI (4’, 6-diamidino-2-phenylindole) fluorochrome applying a modified protocol of (Suda and Trávníček 2006). DNA content measurements of silica dried leaf material was done in a CyFlow Space flow cytometer (Sysmex Partec GmbH, Münster, Germany). Leaf material (about one square centimetre of each sample) was incubated for 80 minutes in 1 ml Otto I buffer (0.1 M citric acid, 0.5% Tween 20) at 4 °C, and then chopped with a razor blade. After incubating for 10 minutes on ice, the homogenate was filtered through a 30 µm nylon mesh. Then the suspension was centrifuged at 12.5 × RPM at 10°C for 5 minutes. This step was repeated two to three times for some samples until pellet of nuclei showed up. The supernatant was discarded and the nuclei re-suspended with 200μL Otto I buffer. Before the samples were analyzed, 800 μL Otto Ⅱ (0.4 M Na2HPO4.12H2O) containing DAPI (4′-6-diamidino-2-phenylindole, 3 μg mL-1) was added and incubated for 30 minutes in the dark to stain the nuclei. The histograms for each sample were evaluated with FloMax V2.0 (Sysmex Partec GmbH, Münster, Germany). *Salix caprea* with known ploidy level (2x=2n=38) was used as an external standard. To re-establish the G1 peak position the standard was used after every tenth sample. The ploidy level was roughly calculated as: sample ploidy = reference ploidy × mean position of the G1 sample peak divided by the mean position of the G1 reference peak.

### Molecular treatment and analyses

The DNA of all samples was extracted using the Qiagen DNeasy Plant Mini Kit following the manufactureŕs instructions (Valencia, CA). After quality check, the DNA was sent to Floragenex, Inc. (Portland, Ore., USA) where the RAD sequencing library preparation was conducted after the protocol described in Baird et al. (2008). The methylation-sensitive restriction enzyme *Pst*I was used for digestion. A former study showed that using this enzyme will cut almost exclusively in the nuclear genome (Gramlich et al. 2018). After size selection of 300bp - 500bp with a Pippin Prep (Sage Science, Beverly, Massachusetts, USA) the libraries were barcoded by individual and multiplexed on an Illumina HiSeq 2500 (Illumina Inc., San Diego, CA, USA). To avoid loss of coverage, we analyzed the diploid and polyploid accessions on two different RAD sequencing plates. The quality of the resulting single-end 100bp long sequence reads was checked using FastQC v.0.10.1 (Andrews 2010).

Sequence reads were de-multiplexed and the fastq files of each sample were used to run ipyrad v.0.7.28 (Eaton and Overcast 2016). The RAD sequencing data were analyzed using a clustering threshold of 85% similarity and a minimum depth of eight reads for base calling. The maximum number of SNPs per locus was set to 20, the maximum number of indels to 8. The adapter trimming option included in ipyrad was used to make sure that all adapters are removed. The consensus sequences of each individual were clustered across samples by sequence similarity of 85%. We set a threshold of maximal four alleles in the final cluster filtering. The resulting clusters represent putative RAD loci shared across samples. Several settings for the minimum number of accessions sharing a locus (m) were initially tested, ranging from m4 (average number of accessions per species) to m120 (loci present in c. 90% of accessions) and optimized as described in Wagner et al. (2018).

The statistics for the tested settings are summarized in Supplement Table S2. Finally, the m40 dataset (29.06% missing data) was used for phylogenetic approaches and the m100 dataset (9.19% missing data) for the genetic structure analysis. The seven steps of the ipyrad pipeline were also performed for each clade that contains one or more polyploid samples to find the optimal number of shared loci and SNPs, respectively, specific for each clade.

We inferred phylogenetic relationships on concatenated alignments of the complete dataset as well as for each clade separately by using maximum likelihood (ML) inferring the GTR+ Γ model of nucleotide substitution implemented in RAXML v.8.2.4 (Stamatakis 2014). Because we used a concatenated data set of short loci, it was not possible to analyse them as separate partitions. However, Leaché et al. (2015) have shown that it is preferable to use full sequence information instead of only SNPs, because this might lead to branch length and accuracy bias.

We performed for each analysis a rapid bootstrapping analysis with 100 replicates using the -f a option, which searches for the best-scoring tree. The resulting trees of all analyses were obtained in FigTree v1.4.3 (Rambaut 2014). Next to the rapid bootstrapping (BS) we used the python script *quartetsampling* (QS; Pease et al. 2018) to infer statistic support for each branch in a given topology. In contrast to bootstrapping, QS is able to distinguish between conflicting signals and poor phylogenetic information. It assesses the confidence, consistency and informativeness of internal tree relationships and the reliability of each terminal branch by using a given topology (in our case the RAxML phylogeny) and the underlying alignment information (=concatenated RAD loci) to build four subsets (=quartets). By sampling randomly samples of each of the four subsets, the likelihood of the concordant topology and the two discordant topologies is evaluated for each branch (see detailed description in Pease et al. (2018)). The first value describes the Quartet Concordance score (QC, [1,-1]), the second the Quartet Differential score (QD, [0,1]) and the third the Quartet Informativeness score (QI, [0,1]). QC gives the support of the current topology, the QD is an indicator of introgression and QI quantifies the informativeness of each branch. The python script (available at https://github.com/FePhyFoFum/quartetsampling) was used with default settings and the obtained alignments and RAxML phylogenies were used as input data. We run 300 replicates using the –L option (minimum likelihood differential). For each phylogeny shown here, the observed QS values (QC/QD/QI) were visualized along with the BS values above and below branches, respectively.

### Species tree estimation using SNAPP

The SNAPP method (Bryant et al. 2012) estimates species trees directly from biallelic markers (e.g., SNP data), and bypasses the necessity of sampling the gene trees at each locus. This is specifically advantageous for RAD sequencing data sets where the calculation of single gene trees is not possible. The method works by estimating the probability of allele frequency change across ancestor/descendent nodes (Bryant et al. 2012). The result is a posterior distribution for the species tree obtained without the estimation of gene trees. For species tree calculation with SNAPP v1.4.2 we used the unlinked SNPs output (one SNP per locus) of the clade-specific ipyrad pipeline, as one prior of the program is the independent nature of SNPs in a data set (see Bryant et al. 2012). We created the input file for SNAPP using BEAUti v2.5.1 (Bouckaert et al. 2014). The multiple accessions of a species were assigned to one taxon. As priors we used default settings for speciation rate (lambda of 1/X), because no prior knowledge existed. We used only variable sites (SNPs), thus non-polymorphic sites were excluded. We run the analysis in BEAST2 (Bouckaert et al. 2014). Because of the immense computational power needed, we limited the number of mcmc generations to 500,000 and sampled every 1000 tree. The resulting trees were summarized in the SNAPP tree analyser tool and finally visualized in FigTree v1.4.3 (Rambaut 2014).

### Genetic structure and network analyses

In order to test for an influence of reticulate evolution on the genetic composition of the included species, especially the polyploids, we explored the genetic structure of each sample for both the complete dataset and each clade separately by using the unlinked SNPs data. They consist of one SNP per locus and represent therefore independent loci. As a side effect, the size of the dataset is decreased and therewith more suitable to the computational power needed by the used program. To avoid bias by too much missing data, we used a dataset composed of loci shared by more than 90% of individuals (m100). The clade specific analyses were performed using the results of the clade specific ipyrad runs. Additionally, we avoided to include samples with unclear species determination or unknown ploidy level in the subsequent analyses. Structure V. 2.3.4 (Pritchard et al. 2000) assigns individuals to ‘populations’ using genotype data and a Bayesian statistic method. Meirmans et al. (2018) described the challenges but also possibilities of using Structure for polyploid samples. They suggest using the recessive allele flag for polyploid samples that will consider genotypic ambiguity that might be present if there are more than two alleles. However, there is no way to enter data from species with different ploidy levels into Structure. Here, we circumvent the challenge of analysing different ploidy levels at the same time. The allelic information of a polyploid sample was summarized to a consensus sequence during the ipyrad pipeline. In the end, diploids as well as polyploids will look the same in the Structure input file (e.g. the four alleles “AATT” will be reduced to “AT”). This way we simplified the complex data and could combine diploids and polyploids in the same analyses. Some information might be lost especially if more than two alleles are originally present (e.g. “AGTT”) or the dosage of alleles is unknown. However, the huge number of informative SNPs that remain in the data set enabled us to analyse the genetic structure. We chose a burn-in of 10,000 and a MCMC of 100,000 replicates, with three iterations of each value of K (K=number of genotypic groups). A higher amount of replications would be preferable; however, the limits in computational power forced us to reduce the number of replicates per iteration. After an initial test run on a smaller dataset, the range of K was set from 2 to 7 in the overall analysis. For the subclades, K was set from 2 up to the number of taxa included. The optimal K value was estimated by inferring the Evanno test revealing the optimal delta K value in Structure Harvester (Earl and vonHoldt 2012).

Reticulate relationships resulting from hybridization or allopolyploidy are not well represented by bifurcating tree topologies (Huson and Bryant 2006; McBreen and Lockhart 2006). In a bifurcating tree topology, the hybrid may be placed in an incorrect sister position to one of the parents or even at a basal branch outside parents, depending on its genomic composition (McDade 1992). Combination of conflicting genomic data sets of hybrids likely results in decreased statistical support for clades and loss of information (Pirie et al. 2009). To overcome this problem, various network methods have been developed to visualize reticulate relationships (Huson and Bryant 2006; Wen et al. 2016). Split-networks are distance-based and represent incompatibilities in a dataset, which reflects reticulate relationships better than tree-building methods (Huson and Bryant 2006). Among the various algorithms for reconstruction of networks, NeighborNet has been most widely used as a clustering method for recognition of species-level relationships (Morrison 2014). Thus, we utilized SplitsTree4 (Huson and Bryant 2006) in order to reconstruct possible network-like evolutionary relationship among the species. Based on the unlinked SNPs data, we generated the split network by implementing NeighbourNet analysis with variance of ordinary least squares. Additionally, a bootstrapping with 1000 replicates as implemented in SplitsTree4 was conducted on the NeighbourNet to test for statistical support of branches. In all analyses, missing data were treated as unknown.

### Hybridization detection

HyDe (<u>Hy</u>bridization <u>De</u>tection) allows testing for hybridization and introgression at a population or species level based on D-Statistics (Blischak et al. 2018a). We used the ‘run_hyde_mp.py’ script to test for putative parent-hybrid combinations with specific respect to the polyploid individuals in our sampling. HyDe tests all possible combinations of input taxa as putative hybrids and parents. It estimates the amount of admixture (γ) and uses p-values to test for the significance of results. While a 50:50 hybrid is characterized by a γ-value of about 0.5, very low levels of admixture (e.g. 0.01 = close to parent P1; 0.99 = close to parent P2) may be indicators for several processes such as ILS and more ancient hybridization. Intermediate γ-values may also indicate older hybridization events that involved the ancestors from the tips in the phylogeny, or recent hybridization events where at least one parent represents a hybrid species. Thus, we used a range of 0.4-0.6 to identify recent hybridization events in the data set, and intermediate ranges of γ= 0.1-0.4 and 0.6-0.9 for older events. We excluded significant values smaller than 0.1 and bigger than 0.9. The complete sequences as well as the unlinked SNP data of the clade-specific analyses were used as input data, as the authors stated this datatype as the ‘most appropriate input’ (Blischak et al. 2018a). We tested two approaches: first, we assigned all individuals of a species to one taxon, and second, we tested all individuals as separate entities. The latter approach is able to better detect individual admixture. HyDe has mainly been applied to datasets based on whole genome shotgun and transcriptome sequencing (Blischak et al. 2018a; Zhang et al. 2019), but it has also been successfully applied to RAD sequencing data (Duran Castillo 2019). To underline the usage of HyDe in RAD sequencing data we performed a preliminary test and included six diploid recently in-situ generated F1 hybrids between *S. helvetica* and *S. purpurea* (Gramlich et al. 2018). The results revealed γ-values of 0.4-0.5 (complete sequence data) and 0.3-0.4 (unlinked SNPs) when combining the true ‘parental species’ (*S. helvetica* and *S. purpurea)* and the F1-hybrids as ‘hybrid’. These values are in accordance with recent studies which showed similar γ-values for modelled and empirical hybrid data (Blischak et al. 2018a; Zhang et al. 2019). Because we were mainly interested in the putative hybrid origin of the polyploid accessions, we screened the results for significant hybridisation events with the polyploids as ‘hybrid’.

### Categorization of SNPs using the SNiPLoid pipeline

The differentiation of different alleles (homolog/paralog/homeolog copies of the same region) is crucial for the interpretation of the observed variation. The ipyrad pipeline conducts a statistic-based paralog filtering to remove paralog regions and allows up to a certain number of different alleles in a final locus, e.g. two alleles for a diploid organism. However, it finally condenses the observed variation within a sample to a consensus sequence. This consensus sequence gives no information on the proportion of the different alleles in a locus. For phylogenetic analyses, the distinguishing variants between the consensus sequences of samples are enough information to reconstruct relationships. However, when working with polyploids, especially in case of allopolyploid origin, the proportion of two or more different subgenomes in the polyploid is of interest. Because of the short sequence length of RAD sequencing reads (and RAD loci), we decided to use a SNP approach rather than a sequence approach to analyze the composition of the polyploid genome.

SNiPloid (Peralta et al. 2013) is a tool developed to analyse SNP composition of recently established allotetraploid crop lineages emerging from diploid parental lineages. Originally, this approach is based on RNA sequencing data. By mapping the RNA reads of an allotetraploid individual to a reference transcriptome of one parent and comparing the observed SNPs and read depth with the second diploid parent, conclusions on the SNP composition of the tetraploid species can be drawn. SNiPloid is an online software tool (http://sniplay.southgreen.fr/cgi-bin/sniploid.cgi) and is able to distinguish five different SNP categories (Peralta et al. 2013, Table 1). In this manuscript we use the same nomenclature as in Peralta et al. (2013): Cat 1/2 = “inter-specific” SNPs means that the observed allele in the polyploid is identical with only one of the parental species. Cat 3/4 = “derived” SNPs is attributed, when the variation observed in the tetraploid is not identified between parental genomes. This hints to a mutation that occurred after the polyploidization event. Finally, cat 5 corresponds to putative homeo-SNPs, i.e., the tetraploid is heterozygous for homeologous alleles of both parental genomes. The remaining portion of SNPs is categorized as “other” and contains SNP combinations that do not fall into the four categories mentioned above.

**Table 1.** Description of the five distinguished SNP categories by SNiPloid. A possible example of the respective allele pattern of the tetraploid in comparison to diploid 1 and diploid 2 is given in the last column. In all other possible occurrences the category “other” is assigned, e.g. if one parental species is heterozygous for a specific site.

| categories | description | diploid 1 | diploid 2 | tetraploid |
| --- | --- | --- | --- | --- |
| Cat 1 | interspecific SNPs | A/A | T/T | A/A |
| Cat 2 | interspecific SNPs | A/A | T/T | T/T |
| Cat 3/4 | derived SNPs | A/A | A/A | A/G |
| Cat 5 | homeo-SNPs (hybrid) | A/A | T/T | A/T |

To use the advantages of this pipeline and to endeavour the SNP composition of the tetraploids in our dataset we adapted the SNiPloid pipeline to RAD sequencing data (Fig. 1). We initially tested a recent diploid hybrid of *S. helvetica* and *S. purpurea* (Gramlich et al. 2018) and revealed 31% Cat 5 SNPs, 6% Cat 1, 11% Cat 2 SNPs, and 23% of Cat 3/4 SNPs, confirming the power of the pipeline. For our study, we utilized a pseudoreference using the sequence information of the putative parental species similar to Chen et al. (2013). Thus, we used the concatenated consensus RAD loci of one putative diploid parental species as reference for each test. Ipyrad generates and stores the observed RAD loci for every single individual. The obtained SAMPLE.consens file was transferred into FASTA format. The indexing of the reference was done in BWA/0.7.12 (Li and Durbin 2009). The sequence dictionary was created with PICARD/2.10.5 using the CreateSequenceDictionary tool (http://broadinstitute.github.io/picard/). The trimmed and filtered reads of the tetraploid species as well as of the second putative diploid parental species were mapped to the indexed reference applying the BWA/0.7.12 mapping tool using the MEM algorithm. The PICARD suite was used to sort the SAM file and to add read groups to the mapped alignment (‘AddOrReplaceReadGroups’). The resulting BAM file was indexed with SAMTOOLS v. 1.8 (Li et al. 2009) and used as input for the Genome Analysis Toolkit (GATK 3.8; McKenna et al. (2010)) to analyze the read depth (‘DepthOfCoverage’) and the observed SNPs (‘HaplotypeCaller’). For the tetraploid samples the --ploidy argument of HaplotypeCaller was set to 4. The generated read depth and the vcf files were eventually used as input for the executable perl script SNiPloid.pl. The results of SNiPloid were written into two tables, one containing the individual SNPs and one containing the summarized SNP count for the five SNP categories mentioned above. SNPs not falling into the specified categories were treated as ‘other’. The results of each tested combination were summarized in a table and pie diagram, given the percentage of each SNP category, and were then compared to each other. Recently established allotetraploid hybrids should show a high proportion of homeo-SNPs, because they contain half of the alleles of diploid ‘parent1’ and the other half of diploid ‘parent2’ and should therefore be heterozygous for most loci. With ongoing speciation, we assume a higher proportion of cat 3/4 SNPs that are specific for the polyploid and originated after the polyploidization event. Backcrossing with one or both parents might lead to an increase of interspecific SNPs (cat 1/2), but these shared SNPs might also be the result of incomplete lineage sorting. The remaining “other” SNPs that cannot be assigned to any category partly represent shared ancient polymorphisms of the whole group.

**Fig 1.**
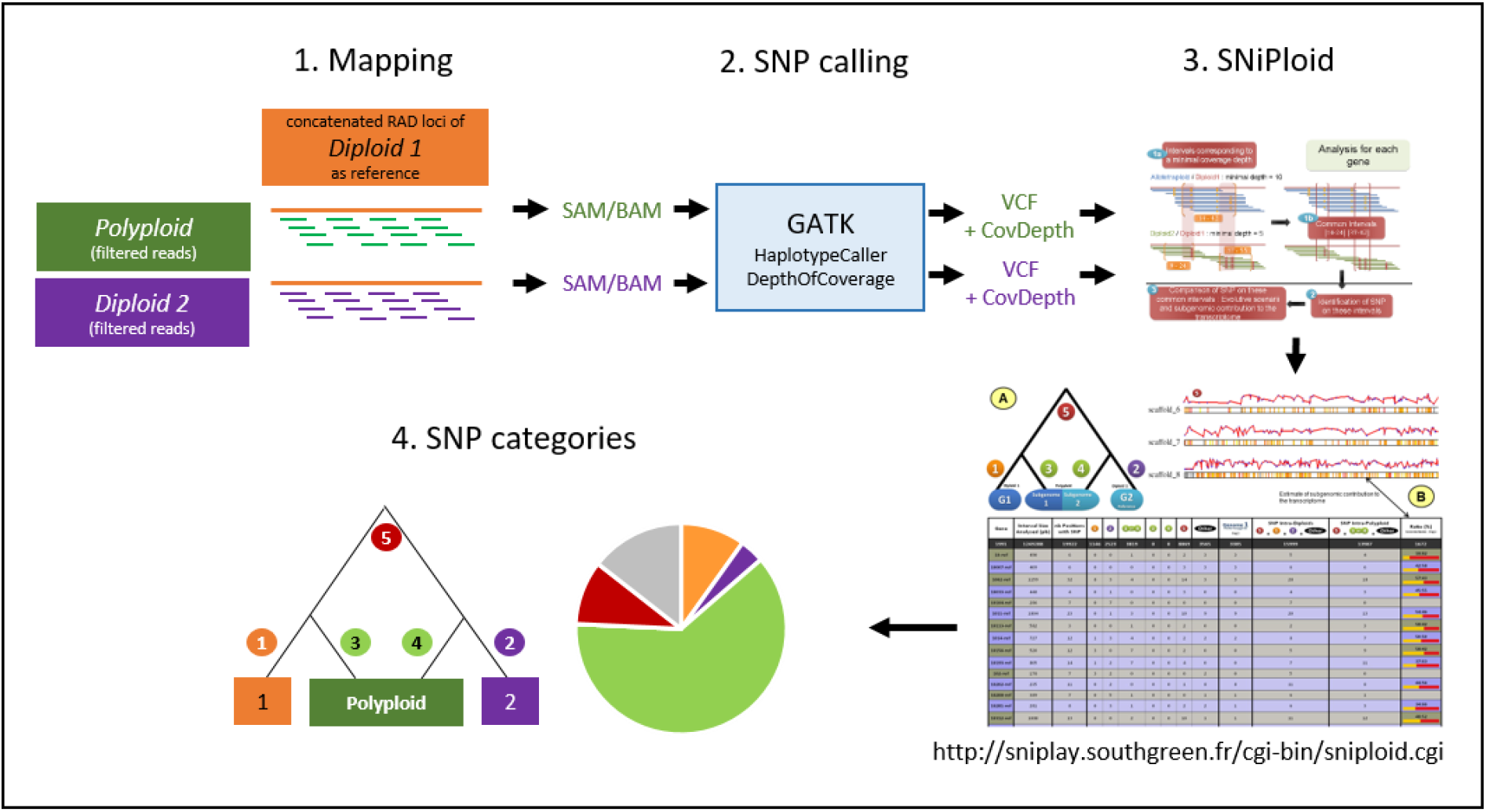
SNiPloid workflow adapted from (Peralta et al. 2013). The filtered RAD sequencing reads of the polyploid and one putative parental species (*Diploid 2*) are mapped to the concatenated RAD loci of the second putative parental species (*Diploid 1*) that act as a ‘pseudo-reference’. SNPs and coverage depth are determined with GATK tools ‘HaplotypeCaller’ and ‘DepthOfCoverage’. The resulting ‘VCF’ and ‘CovDepth’ files of both samples serve as input for SNiPloid. The output of SNiPloid contain the assigned SNP categories for each compared SNP. Those data are finally summarized in a pie chart that shows the proportions of the observed categories.

The pipeline is comparatively time consuming, therefore we did not test all possible combinations of diploid species. Instead, we narrowed down the number of putative parental species combinations by choosing the most likely putative parental species according to the results of the RAxML, genetic structure, NeighborNet and HyDe analyses. Although the extant species might not be the actual parent in the case of ancient hybrid origins, they can be regarded as representative of the parental lineages.

## Results

### Ploidy determination

The sample set included 36 species. For 29 species, we could derive the ploidy level from chromosome counts available in the literature (see Supplement Table S1). To determine the ploidy level of the remaining samples, we analyzed seven species (22 accessions) via flow cytometry. Six of them, S*. integra, S. nummularia, S. pyrenaica, S. pyrolifolia, S. rehderiana*, and *S. schwerinii*, were analyzed for the first time. All analyzed species were diploid (Table S1). Selected histograms are available in Supplement Fig. S1.

### Phylogenetic relationships

The average number of raw reads obtained from the RAD sequencing was 8.15 million reads per sample. The length after removal of adapters and barcode was 86bp. After filtering, an average of 8.08 (+/-5.81) million reads per sample were used for clustering. An average of 161,233 clusters per sample was generated using a clustering threshold of 85% similarity with an average depth of c. 52 reads per cluster. After optimization, we used a data set that consists of 23,393 RAD loci shared by at least 40 individuals and contained 320,010 variable sites of which 191,615 are parsimony informative. The concatenated alignment had a length of 1,931,205 bp with 29.06% missing data (Supplement Table S2). The HyDe results of the m40 dataset revealed 4,724 significant hybridization events of 21,420 tested combination in the taxon-assigned test based on the unlinked SNP data. Of these, 38.6% belonged to combinations that treated a polyploid taxon as ‘hybrid’.

The average γ-value of all observed events was 0.52. The RAxML phylogeny included 133 accessions representing 35 species of the *Chamaetia/Vetrix* clade from Europe and Asia as well as four accessions of *S. triandra* as outgroup (Fig. 2, Supplement Fig. S2). The phylogeny revealed that all species are clearly monophyletic. The topology showed *S. reticulata* in sister position to four main well-supported clades. To keep the results consistent, we used the same Roman numerals for the main clades as in Wagner et al. (2018). Clade I comprised ten species, including two well supported subclades. Clade II contained all members of section Vetrix as well as *S. eleagnos* and *S. serpyllifolia*, clade III comprised all members of sections Villosae and Vimen as well as triploid *S. bicolor*, and clade IV contained *S. hastata*, *S. herbacea*, *S. pyrenaica* and *S. lanata*. The tetraploid species fell into clades I and II, the triploid *S. bicolor* into clade III. The octoploid *S. glaucosericea* was in sister position to clade III and IV while hexaploid *S. glabra* was sister to *S. eleagnos* and *S. serpyllifolia* in clade II, and hexaploid *S. myrsinifolia* was in sister position to the remaining accessions of clade II. The genetic structure analysis of the complete sampling based on 1,655 unlinked SNPs (m100) reflected the four observed main clades (I-IV) and indicated genetic admixture of up to four clusters for the high polyploids *S. glaucosericea* and *S. myrsinifolia* Hexaploid *S. glabra* showed admixture of ‘clade II’ (green) and ‘clade IV’ (yellow, Supplement Fig. S4).

**Fig. 2.**
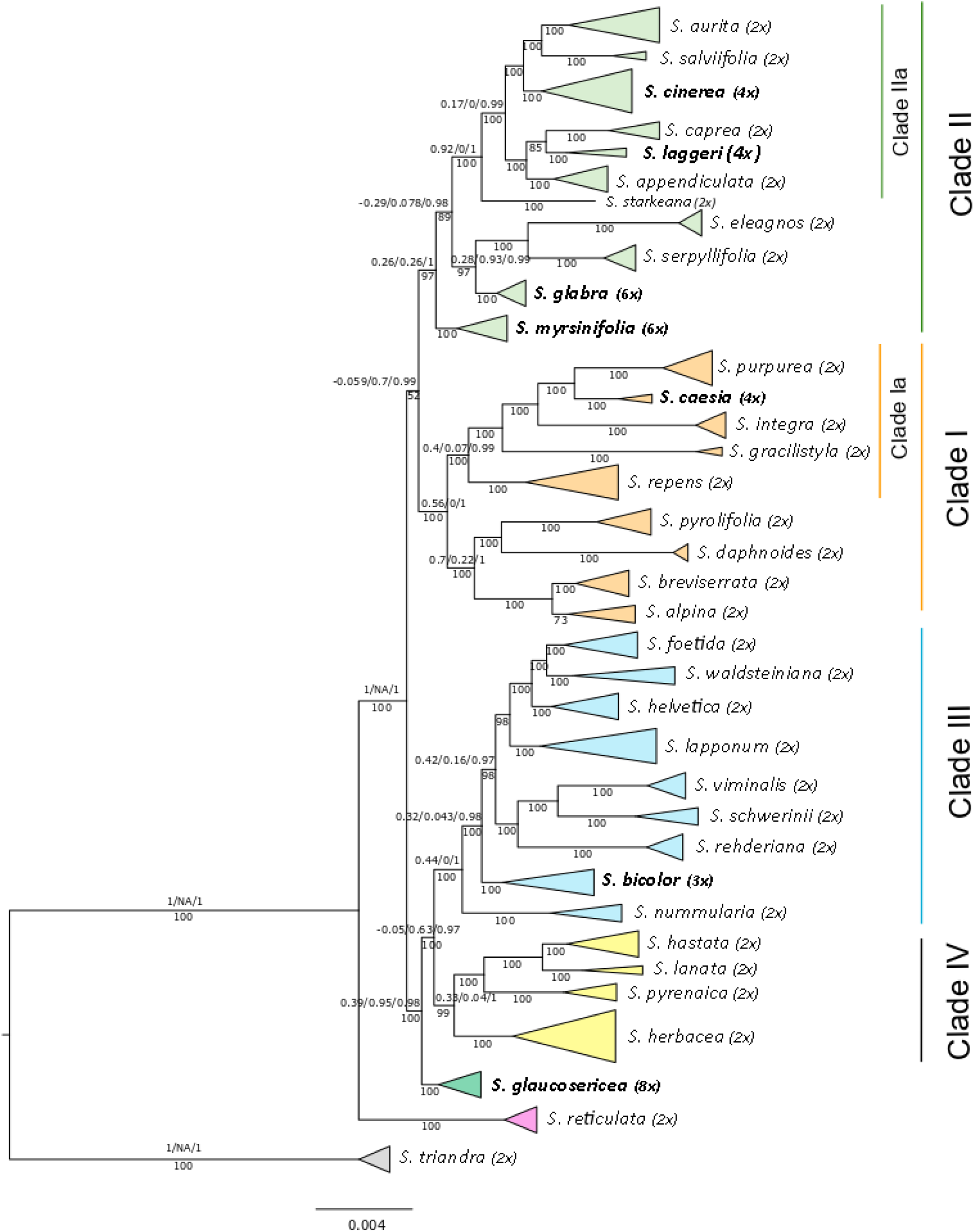
Simplified RAxML phylogeny of 133 accessions representing 36 species of *Salix* subg. *Chamaetia/Vetrix* based on 23,393 RAD loci. Four clades (numbering follows Wagner et al. 2018) are highlighted in different colors, sublclades for detailed analyses are also designated. Ploidy level is indicated behind species name, polyploid samples are highlighted in bold. *Salix triandra* (subg. *Salix*) was used to root the tree. BS values below branches, QS values of selected clades and subclades above branches. A detailed phylogeny including QS support values for all branches is supplied in Supplement Fig. S3.

Each clade that contained one or more polyploid samples was analysed separately to find clade specific loci and SNPs, respectively. The clade specific parameter settings were identical with those of the complete data set and differed only in the minimum number of accessions sharing a locus (m) dependent on the different overall number of accessions in each clade. These were m15 for clade Ia, m23 for clade II and m10 for clade III, respectively.

### Relationships of polyploids in clade I

In the RAxML phylogeny, clade I was divided into two subclades (Fig. 2). The supported monophyletic subclade Ia (BS 100, QS 0.42/0.073/0.99) comprised the species *S. purpurea, S. caesia, S. integra* which belong to S. section Helix sensu Skvortsov (1999), as well as *S. gracilistyla.* They were in in sister position to the closely related species *S. repens* (incl. *S. repens ssp. rosmarinifolia;* sect. Incubaceae sensu Skvortsov (1999). The tetraploid *S. caesia* appeared in a well-supported sister position to *S. purpurea* (BS 100, QS 0.18/0/0.99). The remaining species formed a clade (BS 100, QS 0.7/0.22/1) consisting of two dwarf alpine shrub species, *S. breviserrata* and *S. alpina,* as well as of the shrubs/small trees *S. pyrolifolia* and *S. daphnoides*. The analysis of the 18 accessions of subclade Ia with two accessions of *S. reticulata* as outgroup based on loci shared by at least 15 accessions, contained 38,603 shared RAD loci with 238,614 SNPs. The RAxML topology of the subclade is identical with the complete phylogeny and show strong statistical support for the sister relationship of *S. purpurea* and *S. caesia* (Fig. 3b). The SNAPP analysis of subclade Ia supports the sister position of *S. caesia* and *S. purpurea* in all three observed topologies. In the most abundant topology these two species were in sister position to a clade containing *S. integra* and *S. gracilistyla*. *S. repens* was in an early branching lineage sister to both clades. Using 37,867 unlinked SNPs as input and designating *S. reticulata* as outgroup, HyDe performed a test of 2,448 combinations in the individual approach of which 164 were detected as significant hybridization events. The observed γ-values ranged from 0.2 to 0.9 in 147 events (89,6%), and from 0.4 to 0.6 in 56 hybrid events (34,1%). Only 6 of 30 tested events in the assigned taxa approach were significant. HyDe showed no significant hybridization event with *S. caesia* as hybrid. Thus, we repeated the analysis with the complete sequence data of the clade specific analysis comprising 238,614 SNPs. Here, HyDe detected 606 significant results of 2448 tested combinations in the individual approach with an average γ-value of 0.59. In 85 hybridization events (14%) *S. caesia* was counted as ‘hybrid’. For these events the average observed γ-value was 0.65. The average γ-value was 0.8 in 20 significant combinations with *S. purpurea* and *S. repens* as parental species. Without outgroup the 18 accessions of clade I shared for the m15 dataset 35,078 RAD loci comprising 191,211 SNPs. The genetic structure analysis based on 34,071 unlinked SNPs for the most likely K=4 revealed genetic admixture in *S. caesia* with about 50% of each the genetic partition of *S. purpurea* and *S. repens* (Fig. 3d). In the NeighbourNet analysis (Fig. 3c), *S. caesia* was situated between *S. purpurea* and *S. repens*. To test a hypothesis of allopolyploid origin of *S. caesia* in more detail, we conducted a SNiPLoid analysis using *S. purpurea* and *S. repens* as putative diploid parents of putative allopolyploid *S. caesia*. The results revealed 5.78% Cat 1 and 12.65% Cat 2 interspecific SNPs, 44.52% homeo-SNPs (cat 5) and 19.17% cat 3/4 SNPs. About 17.89% of SNPs did not fall into the given categories and are treated as ‘others’ (Fig. 5b).

**Fig. 3.**
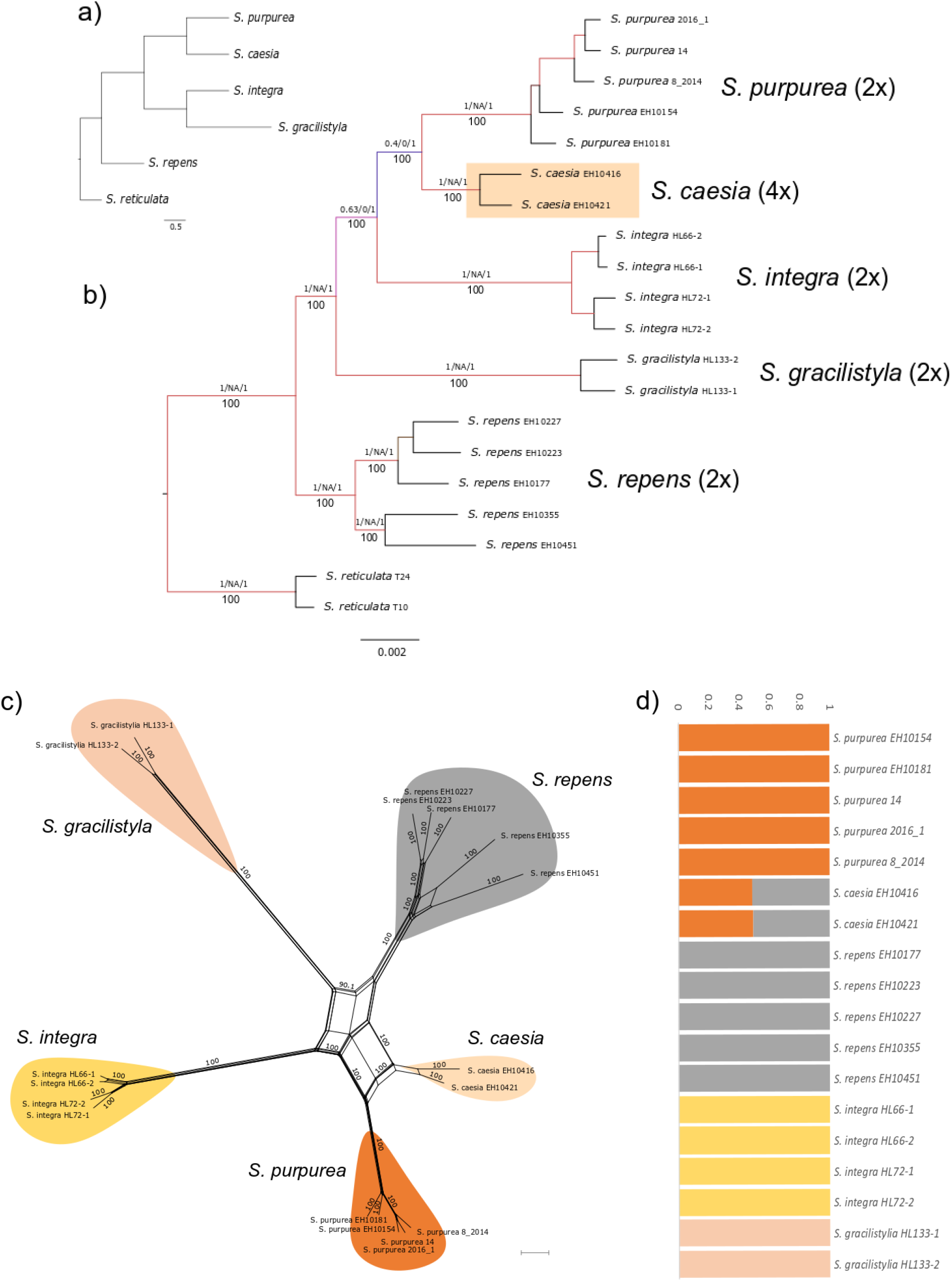
Relationships of tetraploid *S. caesia* in clade Ia. A. RAxML phylogeny based on 38,603 RAD loci and 238,614 SNPs, respectively. Bootstrap support values below, QS values above branches. Genetic structure analysis for the subclade for the most likely value of K=4 based on 37,867 unlinked SNPs. Resulting Splitsgraph of the NeighbourNet analysis of subclade Ia. Bootstrap values indicated at branches.

### Relationships of polyploids in clade II

Clade II contained hexaploid *S. myrsinifolia* and *S. glabra*, both monophyletic. The latter was in sister position to *S. eleagnos* and *S. serpyllifolia*. *Salix myrsinifolia* was in sister position to the remainders of clade II, with good BS but a skew to an alternative topology (BS 97, QS 0.28/0.93/0.99; Fig. 2). All included members of section Vetrix s.l. (sensu Skvortsov (1999)) formed a well-supported monophyletic group (= subclade IIa; BS 100, QS 0.92/0/1, Fig. 2). This subclade contained two tetraploid species, *S. laggeri* and *S. cinerea*. The clade-specific analysis of clade IIa (‘Vetrix-clade’) included 25 accessions and was based on 33,515 RAD loci comprising 254,819 SNPs shared by at least 23 samples (m23). The species were all monophyletic and well supported (Fig. 4b). One accession each of *S. eleagnos* and *S. serpillifolia* served as outgroup. *S. starkeana* was in sister position to the remaining accessions of clade IIa (BS100, 1/NA/1), Tetraploid *S. laggeri* appeared in moderately supported sister position to *S. caprea* (BS100, QS 0.36/0.67/1). The other tetraploid species, *S. cinerea*, was in sisterposition to *S. aurita* and *S. salviifolia* (BS100, QS0.83/0.5/0.99). The backbone of the tree topology showed some less supported branches, especially with regard to the quartetsampling support values that indicate incongruencies of different branching topologies. The most abundant result of the SNAPP analysis (Fig 4a) reflected the QS values by a slightly different topology. *S. cinerea* was in sister position to *S. aurita*, and *S. laggeri* formed a clade with *S. caprea and S. appendiculata* which was also found in the complete dataset (Fig. 2).

**Fig. 4.**
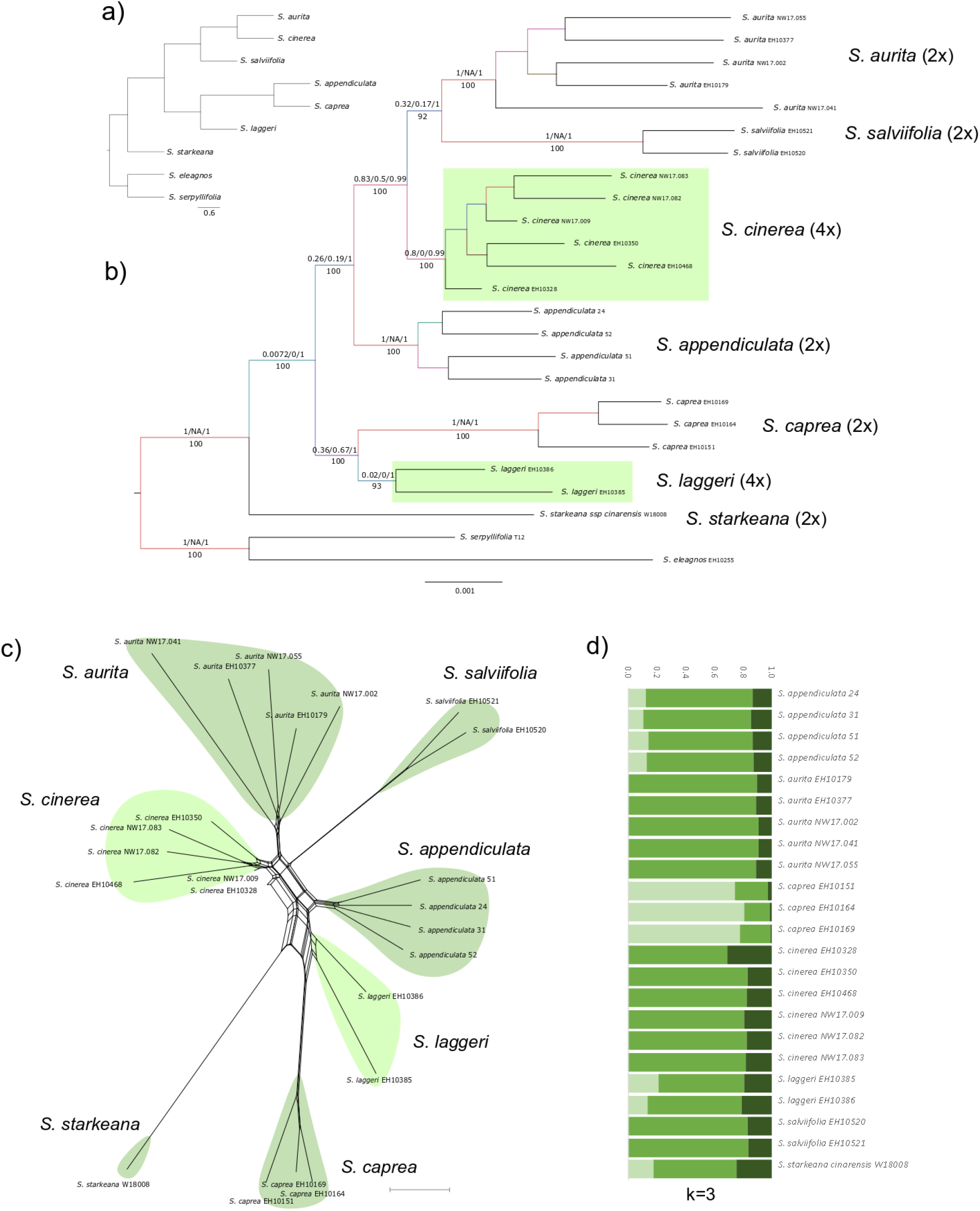
Relationships of polyploids in subclade IIa. RAxML phylogeny based on 33,515 RAD loci, with QS support values above and BS values below branches Tetraploid species indicated with light green boxes. Upper left corner, most abundant SNAPP species tree. Results of the genetic structure analysis for subclade IIa (K=3) based on 39,746 unlinked SNPs, respectively. Splitsgraph of the NeighbourNet analysis of clade IIa, BS values given at branches.

The HyDe analysis of the unlinked SNP data tested 5,313 combinations and revealed 106 significant hybridization events, when we treated the samples as individuals. The taxon-assigned approach tested 104 combinations and yielded only four significant events. In 24 events, *S. cinerea* was treated as ‘hybrid’. The average γ-value for these events was 0.46. Specifically for the parental combination of *S. appendiculata* and *S. aurita,* the nine significant hybrid combinations showed an average γ-value of 0.49. No significant hybrid event was observed for tetraploid *S. laggeri*.

The genetic structure analysis of 39,746 unlinked SNPs without outgroup for the most likely K=3 revealed genetic admixture for all samples. *Salix laggeri* showed an admixed genetic structure comprising mainly the shared partition of the subclade as well as partly the partition of *S. caprea*. *S. cinerea* showed a similar structure as *S. aurita* and *S. salviifolia*. The NeighbourNet based on the same data revealed *S. cinerea* in sister position to *S. aurita* and *S. laggeri* close to *S. caprea* and *S. appendiculata* (Fig. 4c). The tetraploid species *S. cinerea* was situated between *S. aurita* and *S. appendiculata* in the RAxML phylogeny, in sister position to *S. aurita* in the SNAPP species tree and in close relationship to *S. aurita* in the NeighbourNet analysis. This is in accordance with the genetic structure results of subclade IIa that show similar genetic pattern of *S. cinerea* and *S. aurita*. The SNiPloid results for *S. cinerea* using *S. aurita* and *S. appendiculata* as potential diploid parental species, showed about 5.6% homeo-SNPs and 5.6% cat 1 and 5.7% cat 2 SNPs, respectively, whereas about 47.5% were heterozygous sites sharing one allele of only one parent (cat 3/4 SNPs). 35.3 % of SNPs did not fall into the given categories (Fig. 5d).

**Fig. 5.**
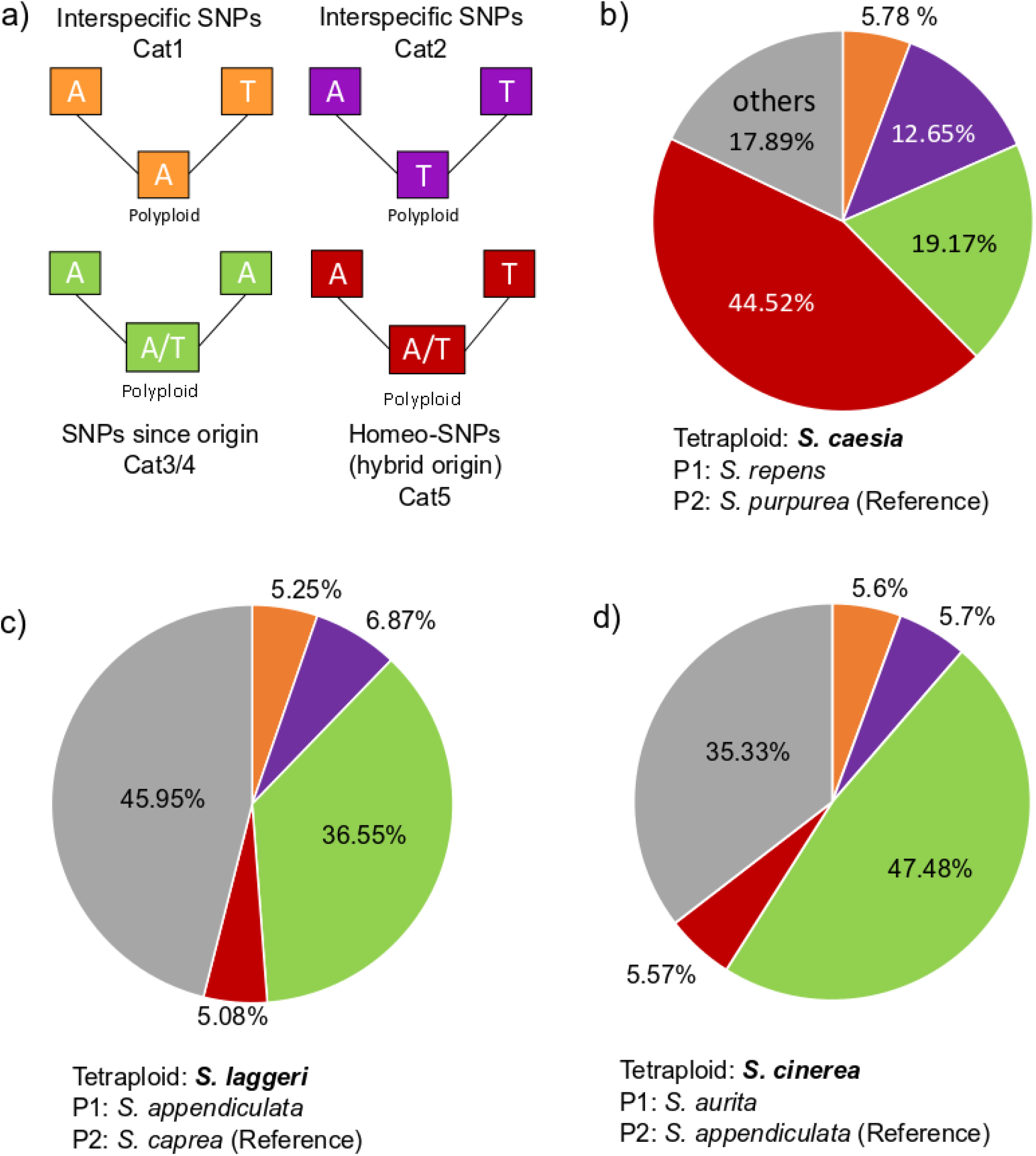
SNiPloid results for the three tetraploid species analysed. *S. caesia*, with *S. repens* and *S. purpurea* as putative parental species, *S. laggeri*, with *S. appendiculata* and *S. caprea* as putative parental species and *S. cinerea* with *S. aurita* and *S. appendiculata* as putative parental species. The colors in pie diagrams represent the proportion of the different observed SNP categories: Cat1, orange, Cat2, lilac (both interspecific SNPs), Cat3/4, green (post-origin SNPs), Cat5, red (homeo-SNPs). Grey indicates the proportion of observed SNPs not falling into the five specified categories. The legend in the upper right corner shows an example of SNP categorization by using the same color code. The parental samples (parent1 and parent2) are diploids. The polyploid SNPs are categorized by comparing the SNP composition with both parents.

*Salix laggeri* was situated in sister position to the remaining species of section Vetrix in the clade-specific analysis. The NeighbourNet showed *S. laggeri* in close relationship to *S. appendiculata* and *S. caprea* (Fig. 4). The genetic structure analysis of subclade IIa for K=3 revealed a similar genetic composition as *S. appendiculata*, with some admixture of the *S. caprea* specific partition (Fig. 4). The SNiPloid analysis with *S. caprea* and *S. appendiculata* as putative diploid parents revealed 5.6% homeoSNPs and about 12.2% interspecific SNPs (5.3% cat 1, 6.9% cat 2). About 36.6% of SNPs were shared with one allele of one parent (cat 3/4), while 45.9% of SNPs were not categorized (Fig. 5c).

### Relationships of polyploids in clade III

Nine species grouped into clade III, which is monophyletic and well supported (BS 98, QS 0.43/0/1). The clade-specific analysis was based on 37 accessions with a minimum of 30 individuals sharing a locus and revealed 16,463 RAD loci containing 97,576 SNPs. *Salix reticulata* was used to root the RAxML tree. As illustrated in Fig. S3, *Salix nummularia* was in sister position to all remaining members of this clade followed by *S. bicolor*, which is triploid and the only polyploid species in this clade. Two subclades diverged: subclade IIIa contained the species *S. helvetica*, *S. waldsteiniana*, *S. foetida*, and *S. lapponum*, while subclade IIIb consisted of *S. viminalis*, *S. schwerinii,* and *S. rehderiana*. Each species is clearly monophyletic. Interestingly, the observed sister relationships of *S. waldsteiniana* and *S. foetida* (S. subsection Arbusculae sensu (Skvortsov 1999) as well as *S. viminalis* and *S. schwerinii* (S. section Vimen sensu Skvortsov (1999)) reflected their respective morphological similarity. These relationships were also supported by neighboring positions in the NeighbourNet analysis (Supplement Fig S3). However, in the clade-specific structure analysis for the most likely K-value of five genetic clusters (Supplement Fig S3), *S. bicolor* showed genetic admixture between *S. nummularia* and a subclade. The amount of admixture differs between the two samples from the Eastern Alps in Austria (about two thirds shared with subclade IIIa) and the two samples from the Harz (Brocken) in Germany (about half shared with subclade IIIa). The deep split between the two geographical regions is also reflected in the NeighbourNet network. SNiPloid was not used to analyse the putative parenthood of *S. bicolor,* for the tool is optimized for tetraploid species (containing two subgenomes) and therefore not suitable for triploid species.

## Discussion

### RAD Sequencing data and analysis pipelines for polyploids

Reduced representation libraries like RAD sequencing are nowadays frequently used to analyze intraspecific population structure, closely related species groups and to infer phylogenetic relationships of diverged lineages (e.g. Cariou et al. 2013; Andrews et al. 2016; Eaton et al. 2017). Especially, when no reference genome is available and a high number of accessions shall be analyzed, reduced representation methods enable researchers to harvest thousands of SNPs at comparatively low costs. However, the use of RAD sequencing for polyploid species is still hampered by the lack of suitable tools, and the statistical difficulties of dealing with more than two alleles (reviewed in Clevenger et al. (2015)). The authors summarized the available analytical tools for SNP calling in polyploids, however, these tools were mainly developed for model plants like crop species, where the parental lineages are previously known. Thus, only few studies using RAD sequencing on polyploid species are published so far (Mastretta-Yanes et al. 2014; Qi et al. 2015; Brandrud et al. 2017, 2019; Feng et al. 2018). Here we present a phylogenomic study on diploid and polyploid willow species by using a de-novo assembly of RAD sequencing data as implemented in ipyrad. Sufficient read depth is important for the quality of SNP calling. Polyploids require an increased depth of coverage based on the bigger genome size and the higher number of alleles (Hirsch and Buell 2013; Clevenger et al. 2015). Thus, we sequenced polyploid and diploid taxa on different plates to avoid loss in coverage. Given that the general minimum read depth per locus in RAD sequencing is six (Eaton 2014), an average depth of c. 52 reads per cluster as observed in our study is enough to endeavor all possible alleles.

The assembly and concatenation of short sequenced fragments representing the whole genome resulted in a robust phylogeny for the *Chamaetia/Vetrix* clade. The method of concatenation has been often criticized, especially in case of conflicting signal among genomic regions. However, Rivers et al. (2016) showed that the data output of the concatenated RAD loci including thousands of unlinked SNPs is able to recover a robust tree. Our findings support the suitability of RAD sequencing for interspecific phylogenetic inference and are in accordance with many studies on different levels of divergence (e.g. Hipp et al. 2014; Eaton et al. 2017; Wagner et al. 2018). The allelic information of the polyploid samples can be condensed to a single consensus sequence to circumvent the challenges of dealing with more than two alleles for phylogenetic approaches. However, the loss of information caused by this simplification might lead to wrong placement of the polyploid accessions in the phylogeny (Eriksson et al. 2018; Andermann et al. 2019). Our results showed that the thousands of informative sites generated with RAD sequencing contains despite this simplification enough information to resolve the phylogenetic relationships. While for phylogenetic approaches complete sequence information is preferable (Leaché et al. 2015), many unlinked loci are the preferable source of information for analyses were the sites are treated as independently evolving genetic entities, e.g. HyDe (Blischak et al. 2018a). The unlinked SNPs (one SNP per locus) of the ipyrad pipeline were also used to reconstruct a species tree using a coalescence based method implemented in SNAPP (Bryant et al. 2012). This method was developed for species tree estimation based on SNP data to circumvent the generation of gene trees. Reduced representation methods generate thousands of biallelic SNPs in short loci that are too small to calculate gene trees. Thus, SNAPP worked well for the SNP data generated with ipyrad in our project. However, since the RAxML analysis already revealed a quite robust tree topology, we used this time-consuming coalescence approach only for the subclade data. Our results confirmed that the coalescent approach using RAD sequencing data is suitable for species delimitation, as recently shown by Brandrud et al. (2019) on *Dactylorhiza*.

To overcome the drawbacks of bifurcating tree topologies for understanding reticulate relationships, we used the informative SNPs to reconstruct networks and to infer genetic structure (Feng et al. 2018). Recent reviews (Dufresne et al. 2014; Meirmans et al. 2018) summarized the problems that appear when frequently used tools that were originally developed for diploids (two alleles) are applied to polyploids. Structure (Pritchard et al. 2000) is generally suited for analyzing polyploids but is problematic when dealing with big data sets, because of increased run time. Other structure methods were especially developed for big datasets, e.g. FastStructure (Raj et al. 2014), but are not able to deal with polyploid data. Here, we decided to use the specific unlinked SNPs output of the ipyrad pipeline for the genetic structure analyses with Structure. Because it is not possible to analyse mixed ploidy levels, we circumvent this by condensing the information of all observed alleles into only two alleles, which is automatically done by ipyrad. We decided to do this, first, because we were able to combine polyploid and diploid accessions this way, and second, allotetraploids do cytologically behave like diploids (Clevenger et al. 2015), resulting mostly in bisomic inheritance. The output comprised information on the genetic composition of the included diploid and polyploid species. The higher polyploids showed genetic admixture, as we expected in case of allopolyploid origin. Additionally, within clade Ia the tetraploid species *S. caesia* comprised two almost equal genetic partitions of putative parents. However, the tetraploid species in the clade-specific analysis of clade IIa did not show a clear pattern of admixture between putative parental taxa. This uncertainty might reflect that the relationships in this widespread and species-rich clade IIa (S. section Vetrix sensu Skvortsov 1999) might be very complex.

We used HyDe (Blischak et al. 2018a) and SNiPloid (Peralta et al. 2013) to reconstruct the genomic constitution of the tetraploid species and to draw conclusions on their origin. HyDe is a tool to detect hybridization in a given dataset. In this study, we used it to identify potential parental taxa for the tetraploid species included. We observed that the number of significant hybridization events depends on the size of the input data. We used the unlinked SNP data in our approach as suggested by Blischak et al. (2018a). The complete alignment using all SNPs, however, revealed a higher amount of significant results. HyDe detected 4,724 significant hybridization events in the complete dataset. About 40% of them were combinations including a polyploid as ‘hybrid’. That means that 60% were indicated as hybridization events between diploid samples. The high amount of natural hybridization in *Salix* is well known, (Skvortsov 1999; Argus 2010), and homoploid hybridization even between distantly related species has been documented (Hardig et al. 2000; Gramlich et al. 2018). However, in this study we observe clearly distinct monophyletic groups for all included species. Additionally, the sampled accessions represent morphologically definite individuals. This is in contrast to the results of HyDe. We assume that the huge amount of underlying data representing the whole genome is an explanation for the observed distinct topology in our analyses, while HyDe uncovers non-visible introgression. Another explanation could be that the amount of data cause significant “false” hybrid combination. However, in this study we focused on the detection of putative parental species of the tetraploids and not on the amount of hybridization in general. Here, we showed that it is possible to use HyDe with RAD sequencing data. We observed a correlation of the number of informative sites we used as input data and the amount of significant results. Therewith we confirm the findings of Blischak et al. (2018a) that more input data reveal results that are more accurate.

SNiPloid was originally developed for transcriptome data (Peralta et al. 2013), however, the application of this tool to RAD sequencing data provided us valuable information about the contribution of putative parental species to the tetraploid genome. In combination with Bayesian genetic structure, HyDe and NeighborNet analyses, we could test for allopolyploid origin and revealed some insights into genome evolution after the polyploidization events. Unfortunately, the tool is only suitable for tetraploid species that consist of not more than two subgenomes (Peralta et al. 2013). However, for RAD sequencing data SNiPloid provides an alternative approach by using biallelic SNPs instead of sequence data to reveal potential parenthood of allotetraploids. This increases the potential to analyse polyploid samples with reduced representation methods. Nevertheless, dealing with polyploid taxa in NGS analyses still require additional specific tools.

### Phylogenetic relationships and origins of polyploids in Salix

The RAD sequencing data revealed a well-resolved phylogeny of the *Chamaetia/Vetrix* clade including 35 Eurasian species plus *S. triandra* as outgroup. Our data are in accordance with other studies that already showed that the subgenera *Chamaetia* and *Vetrix* are no separate entities (Lauron-Moreau et al. 2015; Wu et al. 2015; Wagner et al. 2018), and that dwarf shrubs which were traditionally counted among subgenus *Chamaetia*, evolved several times independently. The present phylogeny included members of all sections sensu Skvortsov (1999) and hence covered the morphological diversity and biogeographical range of Eurasian willows. Our molecular data presented here provide a sufficient phylogenetic framework for reconstruction of the origin of European polyploid species.

All of the 36 included species are monophyletic and the tree topology is well supported. The observed clades are in accordance with a former study on European diploid species by Wagner et al. (2018). *Salix reticulata* is in sister position to all other taxa, confirming the results of former studies (Liu et al. 2016; Wagner et al. 2018) and supporting Skvortsov (1999), who described *S. reticulata* as “the most isolated species of *Chamaetia*”. The samples from Asia fall into the four observed clades.

About 40% of *Salix* species are polyploid (Suda and Argus 1968) ranging from tetraploid to octoploid, rarely to decaploid. The chromosome numbers of the European species, including the polyploids, are well documented (Supplement Table 1), while the ploidy level of the Asian species was partly unknown. Our flow cytometry results showed a diploid level for the six included species that were previously not investigated (Supplement Table 1, Supplement Fig. S1). Although some studies are published about tetraploid species in subgenus *Salix* s.l. (Triest et al. 2000; Triest 2001; Barcaccia et al. 2014) no molecular studies existed so far on the origin of polyploid species of the *Chamaetia/Vetrix* clade. In this study we included seven polyploids from Europe with different ploidy levels to test the suitability of RAD sequencing data for polyploids and to study different scenarios of their evolutionary origin. For the polyploid species included here, no diploid cytotypes have ever been reported, and they all exhibit very distinct morphologies and a composed genetic structure. These features make autopolyploid origins unlikely. The included polyploid species are all monophyletic. However, the species appear scattered over the phylogeny, indicating multiple independent origins of polyploids within the genus resulting from different parental combinations. In so far, *Salix* differs from plant genera in which one allopolyploidization event resulted in post-origin adaptive radiation and speciation, e.g., the Hawaiian Silversword alliance (Seehausen 2004). Willows rather resemble other plant genera with repeated independent polyploidization events, like in *Nicotiana* (Chase et al. 2003), *Achillea* (Guo et al. 2013), *Ranunculus* (Baltisberger and Hörandl 2016), and *Dactylorhiza* (Brandrud et al. 2019), among others.

While the higher polyploid willows are in sister position to major clades or on basal branches, the tetraploids fall within these clades (Fig. 1). This could be due an older origin of the higher polyploids with the contribution of two or more parental lineages that are probably ancestors of the extant major subclades (I-IV), but not conspecific to the extant terminal species. According to the crown group age of the *Chamaetia/Vetrix* clade (23.76 Mill. years, clade B in Wu et al. 2015), their origins may date back to the late Miocene/ early Pliocene. However, without a robust dating of the phylogeny, these estimates remain speculative. The position of a hybrid in a clade can be also influenced by having a parent from a different clade (McDade 1992). Our structure analyses indicated that the hexa- and octopolyploids, but also triploid *S. bicolor*, harbor two to four genetic partitions that characterize otherwise the main subclades I-IV (see Fig. S5). Hence, we assume allopolyploid origins from divergent lineages, potentially also involving more than two parental species in the polyploids with four genetic partitions (*R. myrsinifolia* and in *R. glaucosericea*). However, since the genetic structure analyses cannot disentangle ancient and recent hybridization or introgression events, further conclusions of the origin on the high polyploid willows remain elusive. Hence, we focus here on the putatively younger tetraploids that are situated at the terminal branches of subclades. We wanted to test hypotheses on the putative Eurasian parental species, or, lineages, respectively, that might have contributed to the formation and speciation of the tetraploids.

### Origin and evolution of the allotetraploid species

Since all included tetraploid species are monophyletic and do form well supported clades in our analysis, we do not assume multiple origins with different parental combinations to the tetraploid species. Instead, we assume a single or few hybrid (allopolyploid) origin(s) from a pool of related individuals (Abbott et al. 2013), followed by speciation. We are aware that extant species might not represent the true parental species, and that extinct taxa might have been also involved. Nevertheless, by comparing the amounts of different SNP categories by testing different combinations of putative diploid parents that are the next extant relatives in our observed trees, we analyzed potential parental lineages that might have been involved in allotetraploid species formation. A comparison of the two respective subclades, however, suggests different evolutionary scenarios.

Clade Ia comprised species assigned to S. sect. *Helix* sensu Skvortsov (1999), except for *S. gracilistyla*. However, all these species share the morphological character of connate filaments. The SNAPP species tree revealed a close relationship of *S. caesia* and S*. purpurea*, supporting the RAxML analyses. Based on our genetic structure analyses, tetraploid *S. caesia* combined the genetic partitions that occur in *S. purpurea* and *S. repens*, with about equal proportions, while the two Asian species *S. gracilistyla* and *S. integra* showed each different genetic partitions. The NeighbourNet analysis confirmed these findings and placed *S. caesia* between the *S. purpurea* and *S. repens* (Fig. 3c). These two species are sympatric with *S. caesia* while the two Asian species do not have extant overlapping distributions with the tetraploid (Skvortsov 1999). HyDe did not show any significant hybridization event for *S. caesia* for the unlinked SNP data set, but detected 14% significant combinations based on the complete sequence data. In case of a comparatively young hybridization event we would expect γ-values between 0.4 and 0.6. The high observed average γ-value of 0.8 for the combinations of *S. purpurea* and *S. repens* supports the hypothesis of an older event or other processes like incomplete lineage sorting. In so far, the HyDE analysis corroborates results of our SNiPloid analysis that revealed a high proportion of homeo-SNPs (44.52%) derived from the two parental species, whereas proportions of SNPs from post-origin interspecific events were much lower (18.53% for cat 1+2). However, the observed amount of interspecific SNPs (cat1, cat2) may also be the result of incomplete lineage sorting. Thus, the observed results support the hypothesis of an allotetraploid origin from *S. purpurea* and *S. repens* s.l., with rather few post-origin backcrossing events. Indeed, no extant hybrids of *S. caesia* with the putative parental species have been reported so far. Additionally, hybridization with other species is extremely rare (Hörandl et al. 2012). The considerable proportion of post-origin SNPs (Cat 3/4, 19.17%) specific for the polyploid *S. caesia* indicates an independent evolution of the lineage. Occupation of niches at higher elevations may have contributed to reproductive isolation of *S. caesia* from the two parental species that occur mostly in the lowlands (Hörandl et al. 2012).

The tetraploid species *S. cinerea* and *S. laggeri* are grouping within subclade IIa that comprised all included accessions assigned to section Vetrix sensu Skvortsov (1999). The members of this section share some morphological characters. They are medium sized shrubs with hairy leaves occurring from lowland to montane regions, often in sympatry, and hybridize frequently, also with species outside clade II (Hörandl et al. 2012). A big, shared gene pool that is visible in high proportions of “other”, unassigned SNPs in SNiPloid, as well as high ongoing gene flow may explain that the clade-specific genetic structure analysis revealed no clear species-specific partitions within subclade IIa. However, both *Salix laggeri* and *S. cinerea* showed high proportions of species-specific SNPs (36.6% and 47.5%, respectively), indicating independent evolutionary lineages. The most likely parental combinations for their origin revealed in both cases around 5% of homeo-SNPs derived from hybrid origin, and also low proportions of post-origin hybridization with both parents, supporting allopolyploid origin. We cannot rule out that the true parental lineages were not included here – either due to extinction or to involvement of species that occur nowadays outside Europe (e.g., in adjacent Russia). A long evolutionary history with post-origin hybridization events further affected genomic composition of the hybrids. Interestingly, the two tetraploids differ strongly in the results of HyDe analyses: *S. cinerea* is responsible for 22% of significant hybridization events in the range of a more recent hybrid, while no significant hybrid event including *S. laggeri* was detected. Extant hybridization of *S. cinerea* with other co-occurring lowland species (mainly *S. aurita* and *S. caprea*) may explain this result, while *S. laggeri* is an subalpine species that is reproductively better isolated from the other species and hybridizes only occasionally with the subalpine, sympatric *S. appendiculata* (Lautenschlager-Fleury and Lautenschlager-Fleury 1986; Hörandl et al. 2012).

Our results on tetraploids overall confirmed an allopolyploid origin and a dynamic post-origin evolution of genomes, indicating speciation and evolution of independent lineages. Distribution ranges and ecological niches of the parental species, however, could have fluctuated from the origin of the clade onwards and may have caused various secondary contact hybridizations in different time periods of the Cenozoic (Hewitt 2004, 2011). The relatively high age of the whole *Chamaetia/Vetrix* clade with a crown group age in the late Miocene, and the lack of a dated phylogeny makes it difficult to pinpoint hybridization events to certain geological time periods. According to extant hybridization patterns, isolation of the polyploid willows appears to be strongly influenced by the strength of habitat differentiation. In extant willow species, differentiation along altitudinal gradients appear to be a strong factor preventing hybridization with other species, as discussed by Martini and Paiero (1988), Hörandl et al. (2012) and, Gramlich et al. (2016) for Central European species, and by Huang et al. (2015) on species from Taiwan. These findings support the notion that occupation of a separate niche is important for the establishment of a newly formed polyploid willow lineage.

## Conclusions

Our data demonstrate that high-quality RAD sequencing data are highly informative for the analysis of the origin and relationships of polyploid species. These data allow for the reconstruction of phylogenetic frameworks and give insights into origin and evolution of polyploid species. Although the ipyrad pipeline condenses the underlying information to consensus sequences, it is suitable to resolve phylogenetic relationships on genus level. Based on this simplification, the observed genome-wide variable sites can be directly used for SNP based analyses like genetic structure and NeighbourNet. Additionally, the biallelic SNPs can be used to conduct coalescent-based species tree analyses with SNAPP and hybridization detection using HyDe. However, some information of the allelic diversity might be lost. Here we presented how the concatenated RAD loci can be used as a pseudo-reference. The information of the mapped reads allows inferring information on allele dosage and SNP composition of the polyploid species, as shown here with SNiPloid.

In willows, polyploidization appears to be predominantly connected to hybridization, i.e. to allopolyploid origin, as hypothesized by Skvortsov (1999). These events happened in different time scales in the phylogeny and involved either related species of the same clade, or more putative ancient events involving more distantly related, ancestral lineages. The putative parental taxa, as far as we can resolve them, appear to be plausible in the context of geographical, morphological and ecological patterns. Our data suggest that polyploids harbor considerable proportions of lineage-specific SNPs and managed to establish stable, self-standing evolutionary lineages after their allopolyploid origin. We also found signatures of post-origin hybridization and backcrossing to the parents in the genome structure of the polyploids.Our pipeline combining different analysis tools can disentangle processes influencing genome evolution.

## Supplementary Material

Data available from the Dryad Digital Repository https://doi.org/10.5061/dryad.wh70rxwhx

Supplement Table S1. Studied plant material including detailed sample location and ploidy level

Supplement Table S2. Statistics for ipyrad parameter setting optimization and for the final data set

Supplement Fig. S1. Selected histograms of our flow cytometry results for the tested species

Supplement Fig. S2. Detailed RAxML phylogeny of 133 accessions representing 35 species of *Salix* subg. *Chamaetia/Vetrix* and *S. triandra* as outgroup based on 23,393 RAD loci. Bootstrap support values below, QS values above branches.

Supplement Fig. S3. Relationships of triploid *S. bicolor* in clade III. A. RAxML phylogeny based on 16,463 concatenated RAD loci and 97,576 SNPs, respectively. *S. reticulata* was used as outgroup to root the tree. Bootstrap support values below and QS support values above branches. B. Genetic structure analyses based on 16,104 unlinked SNPs for most likely K=4. C. Splitsgraph of the NeighbourNet analysis of clade III, bootstrap values given at branches.

Supplement Fig. S4. Genetic structure analysis of the complete data set including 133 taxa samples based on 1,655 unlinked SNPs for most likely K=5. Higher polyploids *S. glaucosericea* (8x), *S. myrsinifolia* (6x) and *S. glabra* (6x) are indicated by red boxes.

## Acknowledgements

We thank Jennifer Krüger and Susanne Gramlich for extracting the DNA and support in lab work. We also thank Susanne Gramlich as well as Andrea Danler, Katrin Scheufler, Claudia Pätzold, Jun Zhao, Hai-lei Zheng, Wan-cheng Hou and Gong-ru Zhou for collecting plant material and field assistance.

